# Adaptive Container Service: a New Paradigm for Robust and Optimized Bioinformatics Workflow Deployment in the Cloud

**DOI:** 10.1101/2024.06.25.600641

**Authors:** Kevin Kang, Jinwen Wo, Jon Jiang, Zhong Wang

## Abstract

We propose Adaptive Container Service (ACS), a new paradigm for deploying bioinformatics workflows in cloud computing environments. By encapsulating the entire workflow within a single virtual container, combined with automatic workflow checkpointing and dynamic migration to appropriately scaled containers, ACS-based deployment demonstrates several key advantages over alternative strategies: it enables optimal resource provision to any workflow that comprise of multiple applications with diverse computing needs; it provides protection against application-agnostic out-of-memory (OOM) errors or spot instance interruptions; and it reduces efforts required for workflow development, optimization, and management because it runs workflows with minimal or no code modifications. Proof-of-concept experiments show that ACS avoided both under- and over-provisioning in monolithic single-container deployment. Despite being deployed as a single container, it achieved comparable resource utilization efficiency as optimized Nextflow-managed, multi-modular workflows. Analysis of over 18,000 workflow runs demonstrated that ACS can effectively reduce workflow failures by two-thirds. These findings suggest that ACS frees developers from navigating the complexity of deploying robust workflows and rightsizing compute resources in the cloud, leading to significant reduction in workflow development time and savings in cloud computing costs.

## I. Introduction

Bioinformatics workflows streamline the processing of genomics data using various software tools to gain scientific insights and make significant discoveries [1]. Developing, optimizing, and managing these workflows present many challenges due to both scientific and infrastructural considerations [2]. From an infrastructural perspective, effective workflows often exhibit one or more of the following characteristics: ease of use, ideally being fully automated for production or providing a user-friendly interface for exploratory analysis; reproducibility, ensuring multiple independent runs produce consistent results; robustness, with the ability to recover from unexpected errors related to computing resource availability during execution; and efficiency, with the capability to dynamically deploy more computing resources as data size increases and reduce resources when demand decreases. Balancing these aspects is crucial for the successful implementation and operation of bioinformatics workflows.

The advent of integrated data and analysis systems, such as open-source projects like Galaxy [3, 4, 5] and KBase [6], and commercial platforms like DNAnexus (http://www.dnanexus.com/) and Terra.bio(https://terra.bio/), has simplified the development and execution of common workflows through web user interfaces. These platforms abstract computing resources (high-performance computing or cloud computing), model data projects that integrate data and applications, and provide reproducible and shareable analyses through notebooks. These platforms are particular useful for exploratory analyses. Recent progress in workflow management systems (WMS) like Nextflow (https://github.com/nextflow-io/nextflow) [7], Cromwell (https://github.com/broadinstitute/cromwell), and snakemake [8, 9] are tailored towards robust, automated, and efficient production analyses. Many WMS use domain-specific workflow languages, such as Common Workflow Language (CWL [10]), WDL (https://openwdl.org/), to standardize domain-specific analysis for reproducible results, and some also allow for configuration to customize these analyses.

These systems are specifically designed to tackle the unique challenges in bioinformatics workloads, which often lack robustness and computing efficiency. Many bioinformatics workflows consist of multiple stages, each requiring distinct software and different data types, leading to varied computing resource needs. As shown in Fig 1A, without WMS, one must run the entire workload on a single node or container (Monolithic) with sufficient RAM to meet the needs of the most memory-intensive applications, while also providing enough CPU cores for the most compute-intensive tasks. For example, if one step needs 32 cores and 1GB of RAM, while another step needs 1 core but 100GB of RAM, an instance with at least 32 cores and 100GB of RAM is required. This results in increased costs due to underutilized resources. This mode of deployment leads to conflicting optimization goals: insufficient RAM leads to out-of-memory (OOM) errors, while insufficient CPU cores significantly prolong the workflow’s execution time. In contrast, WMS (Modular deployment in Fig 1B) allocates the appropriate amount of RAM and CPU cores to each application based on its specific requirements, reducing resource waste. WMS can also parallelize the execution of some applications, shortening the overall workflow execution time. When deployed in the cloud, WMS can leverage auto scaling features provided by cloud vendors to adjust computing resources dynamically, according to the workload’s needs. For example, auto scaling group on Amazon Web Services (AWS, https://aws.amazon.com/autoscaling/) can monitor running applications and automatically adjust computing capacity to optimize cloud computing costs. Similarly, Google Cloud Compose2 (https://cloud.google.com/composer/docs/composer-2/optimize-environments#optimization process overview can add or remove computing resources using predefined policies to match a running workload on Google Cloud Platform (GCP). These horizontal scaling strategies (or scaling out) are effective for bioinformatics workloads with parallelized jobs. However, for jobs that run serially on a single node or those that cannot be parallelized, rightsizing becomes challenging.

**Fig. 1.**
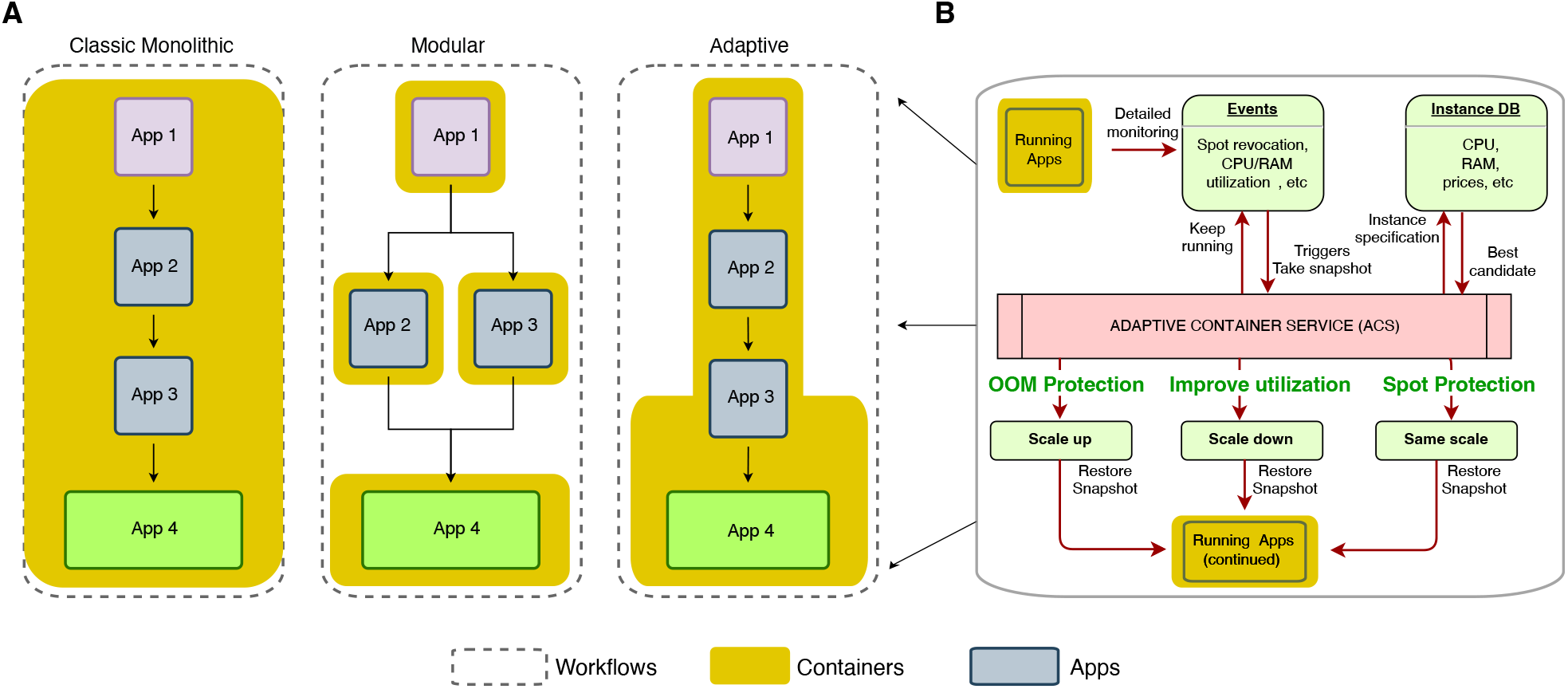
Workflow deployment strategies and implementation of the Adaptive Container Service (ACS). A) In classical Monolithic deployment, the entire workflow, encompassing applications with varied computing resource needs, is encapsulated in a single container (e.g., docker). This container is provisioned with a computational instance in the cloud with sufficient RAM to meet the demands of the most RAM-intensive applications, and an adequate number of CPU cores for the most compute-intensive applications. Insufficient RAM leads to out-of-memory errors, while an inadequate number of CPU cores significantly prolongs the workflow’s execution time. Applications in this setup are executed sequentially. In contrast, in Modular deployment each application is encapsulated in separate containers, and the workflow manager allocates computing instances with the appropriate amount of RAM and CPU cores to match each container based on its specific requirements, therefore enhances resource utilization efficiency. Additionally, some applications can run in parallel (e.g., App 2 and App 3), reducing overall workflow execution time. Monolithic deployment with ACS also executes the entire workflow within a single container. However, the execution on one computing instance can be suspended and dynamically migrated to smaller or larger computing instances based on the resource requirement of the workflow. This migration can also happen in the middle of an application (e.g., App 3). B) The implementation of the Adaptive Container Service. The service continuous monitors container instances for metrics such as CPU and RAM utilization, and instance status. Events like spot instance revocation, out-of-memory (OOM) errors, or prolonged idle periods of computing resources trigger a migration process. The migration process starts with capturing a snapshot of the active container, followed by spawning a new instance, and then restoring the snapshot to it. The type of new instance—larger (scale up), smaller (scale down), or similar (same scale)—is determined by the specific event and the needs of the running apps, as specified in the Instance Database. For example, an OOM error will trigger a scale-up migration, CPU under-utilization will lead to a scale-down, and spot instance revocation will result in a migration to an instance of the same scale.

As workflow management systems (WMS) gain popularity among the bioinformatics community, several limitations become apparent. Learning workflow languages can be challenging due to their steep learning curve. Implementing complex workflows is difficult as the number of applications and their dependencies increase (reviewed in [11, 12, 13]). Each application is packaged in an independent container image, necessitating data exchange through an external file system, which can create overhead in saving and loading data. Additionally, while WMS provide limited robustness by checkpointing intermediate results and allowing the resumption of failed runs, they generally lack granularity in checkpointing. For applications that encounter OOM errors due to unexpected data complexity, manual intervention is often required to change the running environment. In WMS, optimization granularity is typically limited to one step in the workflow, usually corresponding to one application. However, single applications can exhibit large variations in resource utilization during execution, making the complexity of breaking the workflow into more granular steps prohibitively high.

In this paper, we propose a new paradigm for deploying bioinformatics workflows in the cloud to overcome existing challenges. To address the unpredictability and resource wastage in long-running bioinformatics workloads on cloud spot instances, we built a system that monitors workflow execution in real-time and migrates workloads to new containers, leveraging the previously developed Spot-on framework [14], which supports fault-tolerant execution through checkpoint and restart mechanisms. To further protect against OOM errors and resource wastage, we implemented an “adaptive” container service (ACS) that dynamically adjusts the amount of RAM and the number of CPU cores in a running container as needed. ACS actively monitors resource consumption and employs a policy that automatically scales instances up or down on demand. When a workload requires significantly more CPU or RAM, ACS migrates it to a new instance with larger capacity, and vice versa, within a few minutes without losing any work. These innovations allow for continuous workflow execution, optimizes resource utilization, and enhances fault tolerance compared to previous research. We hypothesize that ACS-based workflow deployment can significantly reduce costs and increase the robustness of bioinformatics workloads in cloud environments.

## II. Architecture AND Design

The ACS framework is illustrated in Fig. 1B. For each running application, an agent is responsible for collecting metrics data, including CPU usage, memory usage, network bandwidth, disk usage, and more. This information is transmitted in real-time to the ACS policy engine, where it is analyzed to determine if adjustments to the current container are necessary. The assessment is based on the following considerations:

1. **Real-time Information**: The system prioritizes realtime alerts, such as spot instance eviction events or insufficient system memory, indicating that the application is about to encounter an Out-Of-Memory (OOM) issue. These high-priority problems require immediate attention to prevent workflow interruptions.
2. **Historical Data Analysis**: The ACS policy engine analyzes historical metrics data over a specific period to assess the current operational status. This includes:
  • Checking if the application’s CPU resource utilization aligns with the allocated resources.
  • Ensuring that CPU usage remains above the alert threshold after filtering out temporary peaks.
  • Predicting future memory usage trends to determine if they are likely to exceed the current memory resource limit.
  • Evaluating if disk I/O has reached the device’s upper limit, which could indicate a bottleneck.

These considerations ensure that ACS can dynamically adjust resources, optimizing both performance and cost efficiency while maintaining robust workflow execution.

## III. Algorithm Implementation Details

### A. Computing Resource Monitoring

Continuous monitoring of resource usage for running applications is essential. This includes tracking CPU usage, memory consumption, disk capacity, and network and disk I/O throughput. Based on these continuous monitoring data, our policy engine can determine the running status of the application and assess whether the current instance’s resources are suitable for the application’s requirements.

### B. Workload Migration

Workload migration involves moving a running workload from one machine to another while ensuring continuous operation. From the perspective of the operating system, a running workload consists of one or more running processes/threads. At any given moment, these processes represent a set of states and data across various resources, such as the stack contents of the process, the address space of the process, memory data, and corresponding open files. All these are preserved when an running instance is shut down, and conversely they are restored to a new instance to complete the migration.

### C. Policies for Auto Scaling Up and Down

The ACS policy engine implements strategies for automatically scaling instances up or down based on real-time resource consumption. When a workload requires significantly more CPU or RAM, ACS migrates the workload to a new instance with larger capacity. Conversely, if resource demands decrease, ACS migrates the workload to a smaller instance to optimize resource utilization. These migrations occur within a few minutes, ensuring no work is lost and maintaining continuous workflow execution.

## IV. ACS Enables Optimization OF Single Applications

In the ACS system, computing resource utilization, including RAM, CPU, and I/O, is monitored in real-time. We first tested whether ACS can optimize CPU utilization by dynamically adjusting the total number of CPU cores within a running workflow.

Unlike RAM, incorrectly specifying the total number of CPU cores seldom causes workflow failures. If too few cores are specified, the workflow simply runs for a much longer time. Generally, the running time of a program can be greatly reduced with more cores provided by taking advantage of its multiple processing (MP) capability. However, the number of cores a program can effectively utilize is also affected by factors such as I/O bottlenecks. Provisioning a cloud instance with the appropriate number of CPU cores for an application often requires a deep understanding of the underlying algorithm or the capabilities of the particular instance the application is running on, or both. In both monolithic and modular workflow deployments, the number of cores is often empirically determined, and when historical data is unavailable, a large number may be arbitrarily set. Supplying more cores to an application than it can effectively utilize leads to resource under-utilization.

To demonstrate this situation, we ran the multi-threaded BBTools (sourceforge.net/projects/bbmap/) on a metagenomic dataset in the same instance family (M4 on AWS) with four different numbers of CPU cores and measured the total workflow running time (see Methods for details). Although in theory BBTools can use all the CPU cores of an instance (by default it is set to do so), it only used about 25 of the 40 cores in this experimental setup (Figure 2 ”Over Provision”). Repeating the same experiment with a smaller instance with only 8 cores doubled the total running time from 514 to 1064 seconds (Figure 2 ”Under Provision”). Based on these results, we ran the same experiment with an instance with 16 cores, which took only slightly longer at 626 seconds compared to the 40-core instance (Figure 2 ”Right Provision”). However, the right provision significantly reduced the cost compared to the over-provision case ($0.8/node hour for m4.4xlarge vs $2/node hour for m4.10xlarge on AWS).

**Fig. 2.**
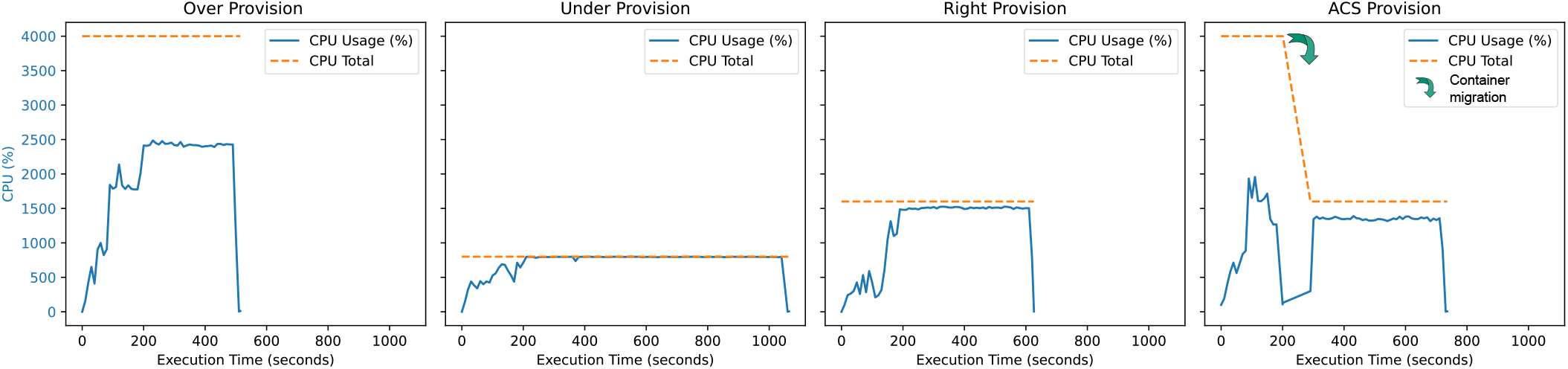
Illustration the concept of ACS optimizes computing resource utilization using a BBTools run as an example. In the case of “**Over Provision**”, only about 25 out of 40 cpu cores were utilized. In “**Under Provision**”, a smaller instance with only 8 cores achieved a 100% cpu utilization but the total running time were much longer. In “**Right Provision**”, an instance 16 cores achieved both 100% cpu utilization and relatively short total running time. In the absence of prior running metrics, “**ACS Provision**” was able to correct an over-provision case and achieved good utilization at a small cost (time spent on migration, green arrow).

In ACS-based deployment, the total number of CPU cores can be dynamically adjusted based on real-time CPU utilization without prior experimentation to determine the right provision. To illustrate this, we repeated the above experiment by first running it on a 40-core instance and then migrating the run to a 16-core instance. The total computing time (646 seconds) was comparable to the right provision case plus approximately 90 seconds for migration (Figure 2 ”ACS Provision”).

## V. ACS Enables Monolithic Deployment TO Achieve Efficiency Comparable TO Modular Deployment WITHOUT Code Modification

Modular workflow deployment, such as with Nextflow, allows for granular, module-specific resource optimization, ensuring robust and efficient workflow execution. Additional measures in workflow management are necessary to coordinate actions across different modules, including the use of a shared file system for data exchange among stateless apps. ACS eliminates these additional measures by running all apps within the same virtual container. Although ACS can also run apps in separate containers in parallel, the following experiment tests the concept of executing an entire workflow in a single container. If we can achieve resource utilization similar to modular deployment, then ACS could enable any monolithic workflow to run efficiently without the complexities of workflow management and optimization.

As a proof of concept, we selected a previously established simple RNA-Seq workflow implemented with Nextflow for testing (Methods). Despite its simplicity, this workflow exhibits common characteristics: it consists of several modules with different computing resource requirements, some steps can be run in parallel, and some steps are dependent on others. We developed a monolithic workflow closely matching the Nextflow implementation (see Data and Code).

As expected, monolithic workflow deployment struggles to balance good computing resource utilization with short running time (Figure 3A). Over-provisioning with 32 CPU cores benefits multi-threaded modules such as Salmon Index and Salmon Quant, but other modules like FastQC and MultiQC, which use only one thread, lead to low resource utilization. Using a smaller instance with only 2 cores resulted in almost 100% CPU utilization but tripled the running time (3 hr 16 min vs. 58 min). Similar to the previous section, ACS provisioning optimized both resource utilization and running time by allocating a large instance for multi-threaded modules and then migrating the workflow to a smaller instance for single-threaded modules (Figure 3A).

**Fig. 3.**
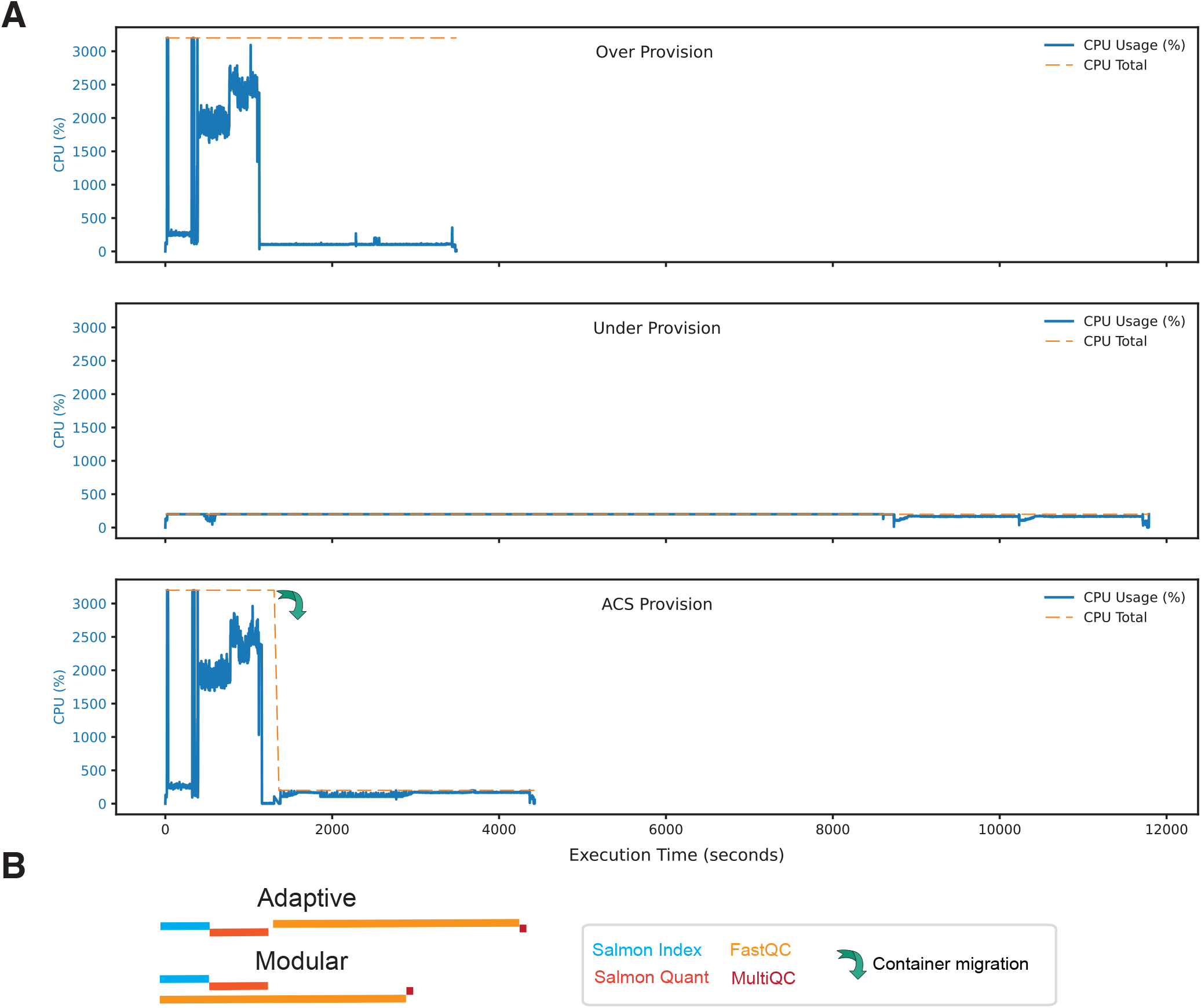
ACS optimizes computing resource utilization for multi-module workflow deployment. (**A**) Monolithic deployment struggles with workflows that include both multi-threaded and single-threaded modules. Provisioning a 32-core instance leads to “**Over Provision**” as CPU cores remain idle most of the time. Conversely, using a smaller instance improves CPU utilization but results in a significantly longer runtime (“**Under Provision**”). With “**ACS Provision**,” a large instance was allocated for multi-threaded tasks, and the workflow was then migrated to a smaller instance (green arrow) for single-threaded modules. (**B**) The comparison between ACS (Adaptive) and Modular deployment shows different colored lines representing various apps. In modular deployment, some modules, such as Salmon Index/Quant and FastQC, run in parallel, while in ACS deployment, they run serially. The total running time is comparable between these two approaches.

The combined running time of all modules between the ACS provision and the modular deployment (Nextflow), which is optimized for resource utilization, is comparable (1 hr 13 min 30 sec vs. 1 hr 3 min 50 sec). However, modular deployment can achieve shorter wall time (53 min 16 sec) if some modules are run in parallel (Figure 3B). It is worth noting that the same exact monolithic code was used for ACS provision without any modification or optimization.

## VI. ACS enables robust workflow deployment on cloud

During the development of ACS, we set up a testing platform on AWS and asked beta users to run common genomics workflows and test its protective ability (See Methods for details). We logged 18,277 independent runs in approximate four months.

We found ACS deployment is able to protect workflow failures due to either spot interruptions and out-of-memory (OOM) errors. Figure 4A shows a short-read alignment workflow by BWA[15] that encountered both OOM (likely due to insufficient RAM provision) and a spot interruption. ACS migrated the workflow to a larger instance (increased total RAM from 72GB to 96GB) to overcome the OOM error, and it migrated the workflow again to a fresh spot instance with the same size when the spot instance it was running on for 10 hours was revoked by AWS. In either case, the workflow would have crashed without running into completion if there were no ACS protection.

**Fig. 4.**
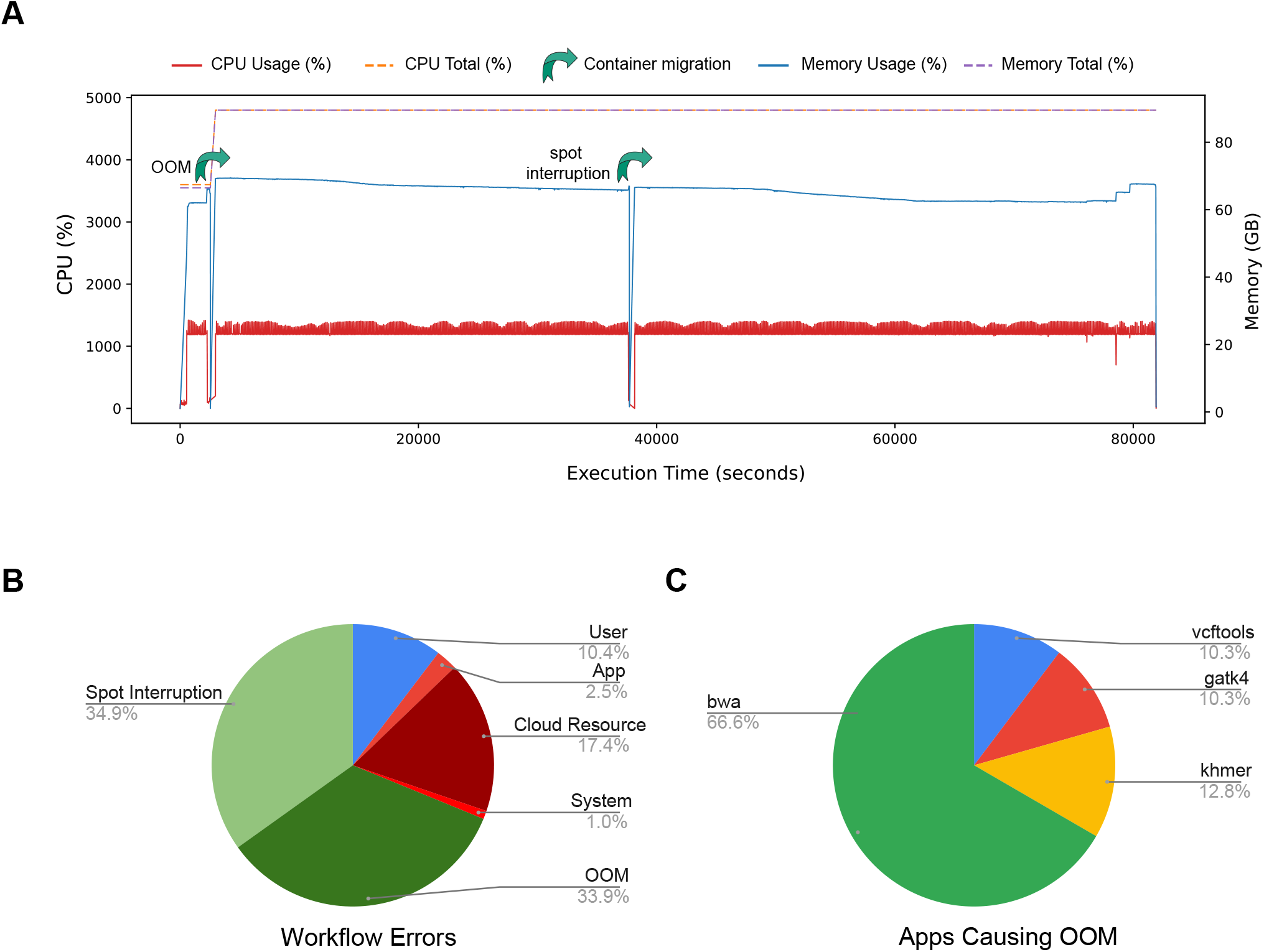
ACS protects workflows from interruptions caused by Out-Of-Memory (OOM) and spot instance interruption on the Amazon Web Services (AWS) platform. (**A**) Metrics for a typical long-running workflow. This is a workload for BWA (Burrows-Wheeler Aligner), a tool used for aligning DNA sequences to a reference genome. The alignment process of BWA mainly involves loading indices, reading sequences, and conducting statistical analysis of insert sizes for paired-end sequences. Depending on the amount of data, these operations require significant amounts of memory and CPU resources. The workflow started on a computation instance with a 36-core CPU and 72GB of RAM. The memory utilization (blue, solid line) quickly surpassed 66GB, approaching the upper limit of the instance’s total RAM (blue, dotted line). The computing job would have crashed because of OOM. ACS detected this risk and promptly migrated the task to a new, larger instance with 48 cores and 96GB of RAM to allow it to continue running. At approximately 10 hour mark (x-axis), the spot instance where the job was running encountered a preemption cloud event that the instance would be reclaimed within two minutes, leading to a forced interruption of the task. Upon receiving the event notice, ACS proactively migrated the running job to a new, same-sized instance as the previous one, and effectively averted a job failure. Eventually the job was successfully completed. Over the entire job execution, total CPU initialization (red, solid line) was relatively low to the total CPU capacity (red, dotted line), suggesting there are other limiting factors than RAM for this particular workflow or insufficient job parameter specifications. (**B**) Among the failed jobs (of running total 18,227 genomics workflow jobs, see details in Methods), the two predominant reasons for job failure are either spot interruptions and OOM errors, accounting for 34.9% (n=256) and 33.9% (n=249), respectively. All other reasons, including user error (User), other types of App crashes (App), cloud resource related errors (Cloud Resource), and failures caused by ACS (System), contributing to less than a third (31.2%, n=229) combined. Spot interruption and OOM errors can be prevented by ACS while the others can not. Errors caused by the ACS system is very rare (1%). (**C**) Common genomics applications that ran into OOM errors (total n=249). The errors were most likely caused by insufficient provision of RAM given the size of the underlying datasets.

Spot interruptions and OOM errors would become predominant workflow errors among the 18000 runs we logged if they are not protected by ACS (Figure 4B). The two accounted for 34.9% (n=256) and 33.9% (n=249) of all workflow failures, respectively. The other failures, caused by user errors (such as software bugs in the workflow), other types of App crashes, or cloud resource related errors, accounted for 30.2%. It is worth-noting that the ACS system in rare cases (1%) can also lead to workflow failures, likely due to snapshot errors.

The protection of OOM by ACS does not depend on specific applications. As shown in Figure 4C, ACS protected several memory-intensive applications, including BWA, Khmer[16], VCFtools[17], and Genome Analysis Toolkit (GATK)[18], from OOM that were likely caused by insufficient RAM, as it is often difficult to estimate the required amount of RAM given a new dataset. Despite the fact that the data presented here is only a small list of potential software tools that can cause OOM, and the percentage of OOM caused by each tool does not reflect the popularity of these tools, these results suggest ACS can effective protect OOM errors and spot interruptions to enable robust workflow runs in an application-independent way.

## VII. Conclusion & Discussions

To enhance the reliability of long-running bioinformatics workflows and improve the utilization of cloud computing resources, we developed the Adaptive Container Service (ACS). ACS-based workflow deployment significantly minimizes Out-of-Memory (OOM) errors and mitigates the risk of spot instance interruptions. This achievement stems from the deployment of an Event Handler that monitors workflow execution for potential interruptions, coupled with a policy engine that migrates the workflow container to different instances to ensure continuous execution. ACS introduces a virtual, elastic computing instance that dynamically adjusts computing resources, offering a novel approach to workflow development and deployment. In comparison to the Monolithic paradigm, ACS maintains workflow development simplicity while enhancing deployment robustness and efficiency. When contrasted with the Modular paradigm, ACS delivers comparable operational efficiency without necessitating extensive development and optimization efforts.

Despite the numerous benefits offered by the Adaptive Container Service (ACS), it is important to acknowledge its limitations. In rare instances (less than 1% of cases), the system may encounter errors stemming from snapshot or migration failures. Additionally, ACS deployment incurs a modest cost, particularly for the storage of snapshots, which could be significant if the workflows requires a large amount of memory. Certain computing instances may also experience limited availability on some cloud platforms, potentially leading to extended wait times or, in some scenarios, migration failures. Moreover, compatibility issues arise with some software tools that do not accommodate dynamic changes in computing resources. These tools, designed to operate with a fixed resource allocation, may continue to limit their resource usage based on initial detection, regardless of subsequent migrations to more capable instances. An example of this is BWA, which only assesses the number of cores at the start of a process and still uses the same number of threads following a migration to a more capable instance with more cores (Figure 4A). For these applications, one workaround is to specify a larger number of threads than the number of cores of the initial computing instance.

The Adaptive Container Service (ACS) may benefit from further enhancement in several areas. A promising avenue for improvement lies in the application of machine learning algorithms to refine resource utilization further. By analyzing patterns and predicting resource needs, machine learning can enable ACS to offer customizable optimization goals tailored to the user’s priorities, such as minimizing total cost or reducing overall execution time. Additionally, we plan to expand the flexibility and efficiency of ACS by providing more options for the elastic computing instance, catering to specific workflow requirements. This includes variations in IOPS (Input/Output Operations Per Second) specifications, which are crucial for optimizing performance in workflows that are heavily dependent on input/output operations.

## VIII. Methods

### A. Running bioinformatics workflows with ACS

To validate the efficacy of the ACS concept, we conducted an experiment using ACS on an internally developed computing platform, dubbed MMCloud (https://www.mmcloud.io/). Over a period of four months, various testing teams actively used this platform to submit a diverse range of computational tasks, totaling 18,277 jobs. Detailed information for each job, including start and end times, the cloud resources consumed during execution, CPU and memory usage, disk load statistics, and any job-related events or errors were captured and are available (see Data and Code availability).

### B. RNA-Seq proof-of-concept workflow

The testing datasets were downloaded from NCBI SRA, and their accession numbers are: SRR24152985, SRR24152986, SRR24152987, SRR24152988, SRR24152989, SRR24152990. The reference transcriptome was GENCODE Human Release 38.

For modular deployment, we used an existing simple RNA-Seq nextflow pipeline (https://github.com/nextflow-io/rnaseq-nf) that consists four modules: fastqc, index, multiQC, and quant. The four modules are carried out by running three software tools: FastQC(http://www.bioinformatics.babraham.ac.uk/projects/fastqc/), Salmon[19], and MultiQC[20]. The nextflow version used was 23.10.1.5891. Steps of Nextflow running on different instances require a shared storage system. In this experiment, we built a distributed file system in the same subnet with a t3.medium instance using the open-source distributed file system, JuiceFS (version 1.1.0+2023-09-04.08c4ae6 (https://github.com/juicedata/juicefs). The Metadata Engine used was in the JuiceFS was Redis (version 6.2.7) and the Data Storage used was AWS S3.

To build a matching monolithic/ACS deployment, we first developed a bash script using the exact program and parameters as the nextflow pipeline. The script was then encapsulated in a customized built docker container for monolithic/ACS deployment.

All experiments were done on AWS using spot instances. Over-provision was done on a c5a.8xlarge instance with 32 cores and 64GB RAM, while under-provision on a c5a.large instance with 2 cores and 4GB RAM. ACS provision was initiated on a c5a.8xlarge instance, and subsequently migrated to a c5a.large instance.

### C. Resource provisions for BBtools

BBtools v39.06 was provided in a docker container (docker.io/bryce911/bbtools). bbduk was run to remove sequence noise using the command “bbduk.sh in=read1 out=filtered1 k=23 ktrim=r mink=12 hdist=1 minlength=50 tpe tbo” on a dataset from CAMI2 synthetic dataset, specifically 3 samples from rhimgCAMI2 short read dataset [21]. All experiments were done on AWS using spot instances and MemVerge’s MMCloud v2.4.0 for over-provision (m4.10xlarge,40 cores, 160GB RAM), under-provision (m4.2xlarge, 8 cores, 32GB RAM), and right-provision (m4.4xlarge, 16 cores, 64GB RAM). ACS provision was initiated on a m4.10xlarge instance, then was migrated to a m4.4xlarge instance.

### D. Code and data availability

All job running data (Figure 2-4) can be downloaded at Zenodo: DOI: 10.5281/zenodo.12533516. Monolithic/ACS RNA-Seq pipeline is available at: GitHub (https://github.com/MemVerge/acs)

## IX. Glossaries

OOM: Out-of-Memory
ACS: Adaptive Container Service
AWS: Amazon Web Service
Container: A lightweight, stand-alone, and executable software package that includes everything needed to run a workflow, including both the software libraries and their settings. Containers are isolated from each other and the host system, providing a consistent environment across various stages of workflow development and deployment.
Instance: A virtual server that runs applications and containers in a cloud computing environment. Instances can vary in size and capacity, and they can be created, managed, and terminated as needed.
Spot Instance: A type of cloud computing instance offered at a lower price than on-demand instances, utilizing spare capacity. Spot instances can be reclaimed by the cloud provider when the resources are needed elsewhere, which can lead to workflow interruptions if unmanaged.
IOPS: Input/Output Operations Per Second, a performance measurement used to benchmark the speed at which storage devices, such as hard drives and solid-state drives, can read and write data.

## XI. Author Contributions

J.Jiang and K.Kang conceived the project. K.Kang and Z.Wang designed the experiments. K.Kang and J.Wo carried out data analyses. All authors authored, reviewed, and approved the final draft.

## XII. Acknowledgements

The authors thank Drs. Charles Fan and Cedric Druce, Mr. Bernie Wu, and Mr. Beau Beauchamp for discussion and critically reviewing the manuscript.

